# Colour Vision in Stomatopod Crustaceans: more questions than answers

**DOI:** 10.1101/2021.10.21.465241

**Authors:** Amy Streets, Hayley England, Justin Marshall

**Affiliations:** Queensland Brain Institute, University of Queensland

**Author notes:** Corresponding author: Amy Streets.

**Keywords:** colour, vision, stomatopod, behaviour

## Abstract

Stomatopod crustaceans, or mantis shrimps, are known for their extensive range of spectral sensitivities but relatively poor spectral discrimination. Instead of the colour-opponent mechanism of other colour vision systems, the 12 narrow-band colour channels they possess may underlie a different method of colour processing. We investigated one hypothesis, in which the photoreceptors are proposed to act as individual wave-band detectors, interpreting colour as a parallel pattern of photoreceptor activation, rather than a ratiometric comparison of individual signals. This different form of colour detection has been used to explain previous behavioural tests in which low saturation blue was not discriminated from grey potentially because of similar activation patterns. Results here, however, indicate that the stomatopod, *Haptosquilla trispinosa* was able to easily distinguish several colours, including blue of both high and low saturation, from greys. The animals did show a decrease in performance over time in an artificially lit environment, indicating plasticity in colour discrimination ability. This rapid plasticity, most likely the result of a change in opsin (visual pigment) expression, has now been noted in several animal lineages (both invertebrate and vertebrate) and is a factor we suggest needing care and potential re-examination in any colour-based behavioural tests. As for stomatopods, it remains unclear why they achieve poor colour discrimination using the most comprehensive set of spectral sensitivities in the animal kingdom and also what form of colour processing they may utilise.

## Introduction

Mantis shrimp, or stomatopods, possess perhaps the most complex retina of all visual systems known (Marshall, 1988; Cronin and Marshall, 1989; Marshall et al., 2007; Marshall and Arikawa, 2014). With 12 spectral photoreceptors (and others for polarisation and intensity detection bringing the total of input-channels to 20), they outnumber, with the possible exception of butterflies (Arikawa, 2003; Chen et al., 2016; Marshall and Arikawa, 2014), the receptor diversity of other animals, which commonly have between two and four spectral sensitivities (Barlow, 1982; Kelber and Osorio, 2010). The 12 colour receptors are spread evenly through the spectrum sampling from just below 300 nm to above 700 nm but most likely do not construct the dodecahedral colour space they are capable of, as there are no known colour tasks in nature requiring this degree of scrutiny (Barlow, 1982; Kelber and Osorio, 2010; Marshall and Arikawa, 2014). Their sharply tuned photoreceptor set has been proposed as effective in achieving colour constancy in the spectrally challenging marine environment (Osorio et al., 1997), but this idea remains hypothetical.

While early behavioural tests demonstrated colour vision in stomatopods based on the von Frisch colour from greys paradigm (von Frisch, 1974; Marshall et al., 1996), more recent and more detailed wavelength discrimination experiments suggest that stomatopods lack fine spectral discrimination (Thoen et al., 2014). Based largely on anatomical evidence (Marshall, 1988; Marshall et al., 1991a,b) a four-spectral window opponent comparison was originally proposed in which different eye regions (rows of ommatidia in the mid-band region of the eye) analysed discrete zones of the 400-700nm spectrum. This would still enable very fine spectral analysis, in particular due to the sharp sensitivities the eye achieves with serial filtering mechanisms (Marshall, 1988; Cronin and Marshall, 1989). Contrary to this, the results of Thoen and colleagues clearly showed that, at least in a two-choice food reward paradigm, spectral discrimination, or Δλ, was unexpectedly poor, indeed around ten times worse than other animals tested in similar ways, including goldfish, butterflies, birds and humans (Thoen et al., 2014; **Figure 1B**).

**Figure 1:**
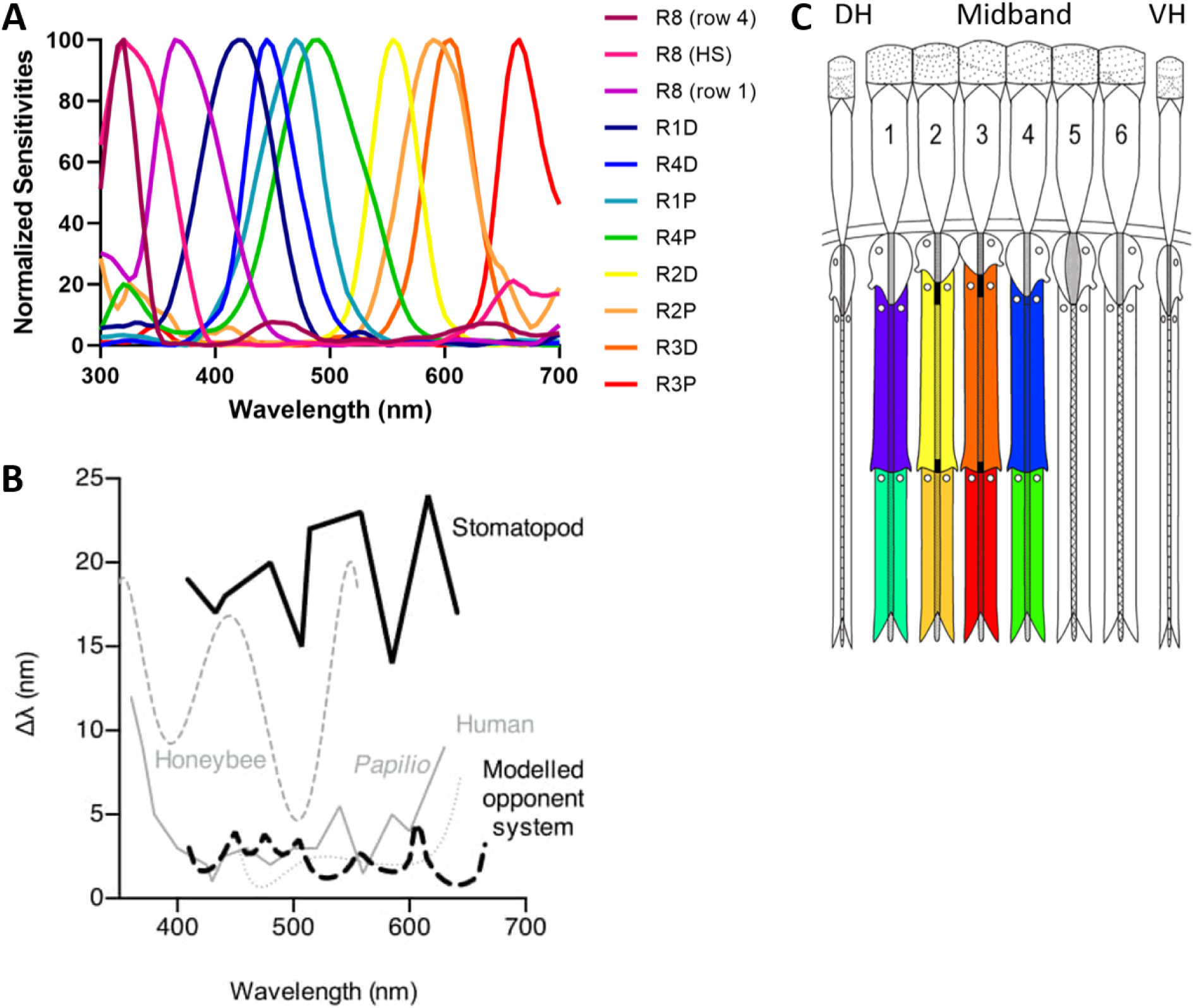
Model of stomatopod (*H. trispinosa*) visual system. **A)** Normalized electrophysiological response of each type of photoreceptor in first 4 midband rows. **B)** Spectral discrimination ability at different wavelengths for several animals. See text for discussion and reference. **C)** Stylized midband showing approximate photoreceptor type sensitivity as colours. The distally placed UV sensitive R8 cells are not coloured. Numbers refer to the respective midband rows and the colours the approximate spectral sensitivity of each tier. **A** and **B** are from Thoen et al., 2014; **C** adapted from Marshall et al., 2007.

Although surprising, these results did potentially provide an explanation for one of the observations made in the original colours from grey behavioural assay. In this experiment, the peacock mantis shrimp, *Odontodactylus scyllarus*, could not discriminate a light blue feeding container from greys (Marshall et al., 1996). In an attempt to explain this failure in choice, it was suggested that mantis shrimp may analyse colour as a pattern of 12 excitations across the spectrum, rather than with any comparison of spectral sensitivities. This idea is congruent with their scanning eye-movements and way of examining coloured objects (Land et al., 1990; Land, 1999).

The reason stomatopods may have evolved such an apparently different visual system is unknown, but evolutionarily they have been out on a limb for 400 million years (Ahyong and Harling, 2000), hinting that they have been ‘trying out’ something different to other animals for a while. It is possible they have a neural mechanism that interprets colour somehow very differently to the typical opponent systems (Thoen et al., 2014) but current investigations into their sub-retinal wiring and potential colour-information processing, at least at the anatomical level, seems to argue against this (Kleinlogel and Marshall, 2005; Templin, 2017; Streets, 2021). There is no totally novel reorganisation in basic neural patterns beneath individual photoreceptors at the lamina, medulla and lobula stages of information processing, either between eye regions or indeed in comparison to other arthropods (Strausfeld and Andrew, 2011; Thoen et al., 2017, 2018).

Stomatopod compound eyes are composed of two peripheral hemisphere regions on either side of a central midband. While the morphology of the ommatidia in the hemispheres is much like other crustaceans, the six rows of ommatidia in the midband are modified in a number of ways. It is in the top four midband rows that the specialisations for colour vision are found, each row sensitive to three wavelength zones. The colour photoreceptors are sharpened and shifted in their spectral sensitivity by a number of filtering mechanisms including short wavelength filtering (in the UV) by the dioptric crystalline cone elements, photoreceptor tiering and, in rows 2 and 3, photostable colour filters (Bok et al., 2014; Cronin and Marshall, 1989; Marshall, 1988; Marshall et al., 2007) (**Fig. 1A,C**). Beneath the retina, information from the midband rows initially remains separated from the hemispheres, and through the eye-stalk neuropils, lamina, medulla and lobula, there are anatomically segregated zones that receive input from midband and each of the hemispheres.

While, as noted already, the basic arrangement here appears similar to other arthropods, in fact centripetally there is increasing complexity and some cross-talk between retinal zones, including between the midband rows themselves (Thoen et al., 2018). Until effective electrophysiological recordings are made at these levels, further speculation is just that. Nonetheless, a brief review of past ideas, placed in the context of knowledge level of the time, contributes to the background and motivation for the behavioural data presented here.

There are several hypotheses previously suggested to explain how mantis shrimps may process colour information. Based on initial anatomical evidence, it was originally proposed that photoreceptor signals outside the ultraviolet (UV) range, may be compared within each of the four colour-sensitive midband rows (1-4) (**Fig. 1C**), delivering a possible “dichromatic” opponent system, each row examining a limited spectral zone from 400-700 nm. Compelling evidence here is in the fact that the rhabdomeric cells in each tier of these rows are the same as those that, in other crustaceans and in the stomatopod hemispheres, are set up as polarisation opponent sub-populations, comparing, for example, horizontally polarised light to vertical (Glantz and Miller, 2002; Marshall et al 1991a; Strausfeld and Nassel, 1981). This means that, without the need to reorganise existing wiring beneath the retina, a polarisation opponency becomes colour opponency.

The UV sensitive rhabdomeric cells, called R8 by convention, in common with many arthropods, possess long visual fibre axonal connection that penetrates through the lamina to terminate in the medulla. In order for any form of intra-UV opponency to be possible (a possibility as supported by the recent work of Bok et al., 2018), an inter-row comparison must be made between R8s (Kleinlogel and Marshall, 2005; Marshall, 1988; Marshall et al., 2007). UV sensitivity in many other animals seems designated to a specific task, such as phototaxis towards UV in flying insects, as it typically represents open sky (Baden and Osorio, 2019; Kelber et al., 2003; Schnaitmann et al., 2020). While this may also be the case for stomatopods (Bok et al., 2018), there is potential for the more complex intra-UV opponency, as they are equipped with 4 separate UV visual sensitivities, peaking from as low as 315 nm. As previously noted, recent behavioural tests support this intra-UV opponency hypothesis (Bok et al., 2018).

An alternative idea is that stomatopods may analyse colour information in a manner similar to the processing of auditory information by the cochlea, examining the chromaticity of light within each spectral band as a continuum of frequency, rather than using opponent processing (Marshall et al., 2007). Instead of comparing spectral signals, downstream computation centres would determine colours by the pattern, or placement in the frequency continuum, of photoreceptor activation. This idea, sometimes called the ‘bar-code’ hypothesis, likening the scan over objects to bar-code readers or indeed QR-codes. The idea has been used to explain the limited colour discrimination ability and argue for a system in some ways more like a colour categorising system based on which photoreceptors are activated. Supporting evidence here comes in two forms. Firstly, there are striking similarities to the way colours are processed in the inferior temporal cortex of primates (Zaidi et al., 2014) and may facilitate faster processing at the cost of poorer colour discrimination between similar wavelengths (Thoen et al., 2014). Secondly, the compressed optics of the stomatopod eye (Marshall and Land, 1993) drive the system to sample the world with slow scanning eye movements (Land et al., 1990). The stomatopod eye therefore can be thought of as operating like a satellite-born push-broom detector or indeed any line-scan device such as a photocopier (Wolpert, 2011).

The bar-code hypothesis also explains the apparent over-proliferation of spectral sensitivities and their narrow-band tuning, as this set of 12 is needed to cover the available spectrum from 300-700 nm (Marshall and Arikawa, 2014). As shown by Barlow and others (Barlow, 1982), an opponent system only requires around 4 spectral sensitivities over this range to decode almost all colour information present on earth.

There is in fact no reason why both opponency and some sort of pattern, barcode, categorical of frequency analysis system might not operate simultaneously. If stomatopods have divided up the spectrum into dichromatic bins, it may be that each of these is used to solve some sort of relevant task while an overall sense of colour is provided by bar-code scanning. Having, in some ways an over-precise opponent process, has been suggested as a way to solve the problems of colour constancy under water (Osorio et al., 1997). In order to further explore the various hypotheses, we conducted a series of experiments using a different species of stomatopod, *Haptosquilla trispinosa*, to determine whether all low saturation colours, not just blue (Marshall et al., 1996), were more difficult to learn than high saturation colours (Experiment 1). Based on initial results, we also set out to examine any inconsistencies in performance, such as a change in visual performance after being kept in captivity (Experiment 2). Potential species difference led us to examine innate preferences for specific colours or colour types (Experiment 3).

## Methods

### Animal Care

Stomatopods were collected on the shallow reefs around Lizard Island Research Station (LIRS) (GBRMPA Permit no. G17/38160.1) during August 2018, 2019, and 2020. They were individually housed in aquaria with small PVC tubes to use as burrows and fed small pieces of raw shrimp. Experiments requiring training were conducted, either at the Lizard Island research Station, under shaded but natural daylight, or at the University of Queensland (UQ), on a 12:12 h light:dark cycle with salinity between 32-35 ppm. The UQ aquarium lights consist of fluorescent tubes combined to provide illumination as close as possible to natural daylight (**Fig. 5**).

### Stimulus Design

The stimuli were made of small white cable ties (2.5mm width, 10cm length), the ratchet-end of the cable tie making a convenient small feeding dish and flat end on which to attach various coloured or neutral density filters (Lee Filters, Andover, UK; see Templin, 2017 for further details). Filters were: high and low saturation red (Lee Filters 182/035, respectively), orange (287/162), green (124/725), blue (195/725), and neutral density filters (Lee Filters: ND 0, 0.15, 0.6, 0.9). The low-saturation blue filter (725) is the same one used in Marshall et al. (1996). ND filters were also the same in Marshall et al. (1996) and other experiments (review: Kelber et al., 2003) to ensure the animals made choice based on colour, not brightness.

**Fig. 2** shows spectral measurements of each stimulus filter attached to a cable tie. In order to make relevant comparisons, spectra were normalised to the maximum reflectance of the transparent filter ND 0. Photoreceptor responses were calculated using a modified quantum catch calculation (Kelber et al., 2003; Marshall et al., 1996). Individual spectral sensitivities for *H. trispinosa* were taken from Thoen et al., 2014 (**Fig. 1A**) and downwelling light measured with an Ocean Optics USB2000 spectrophotometer (**Fig. 5**, “Experiment 1”). These values were normalised by the individual spectra of the filters (**Fig. 2**) to give the final photoreceptor response (**Fig. 3**) as per Marshall and Vorobyev (2003).

**Figure 2:**
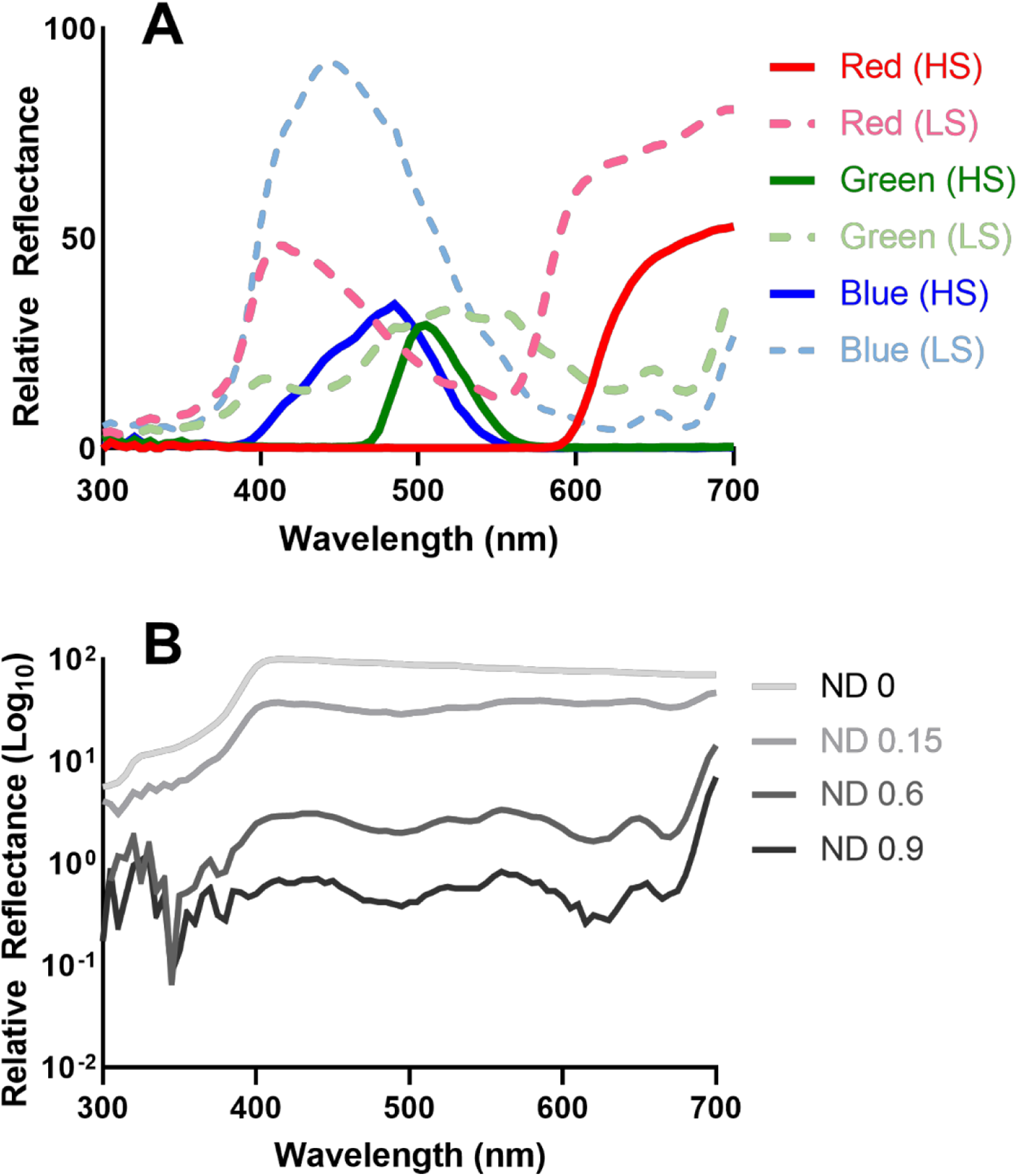
Spectral reflectance curves of stimulus filters. Spectral reflectance measurements of **A)** coloured filters and **B)** neutral density filters. All curves have been normalized to ND 0 in order to demonstrate relative brightness. Highly saturated (HS) coloured filters are solid lines, and less saturated (LS) coloured filters are dashed in **A**.

**Figure 3:**
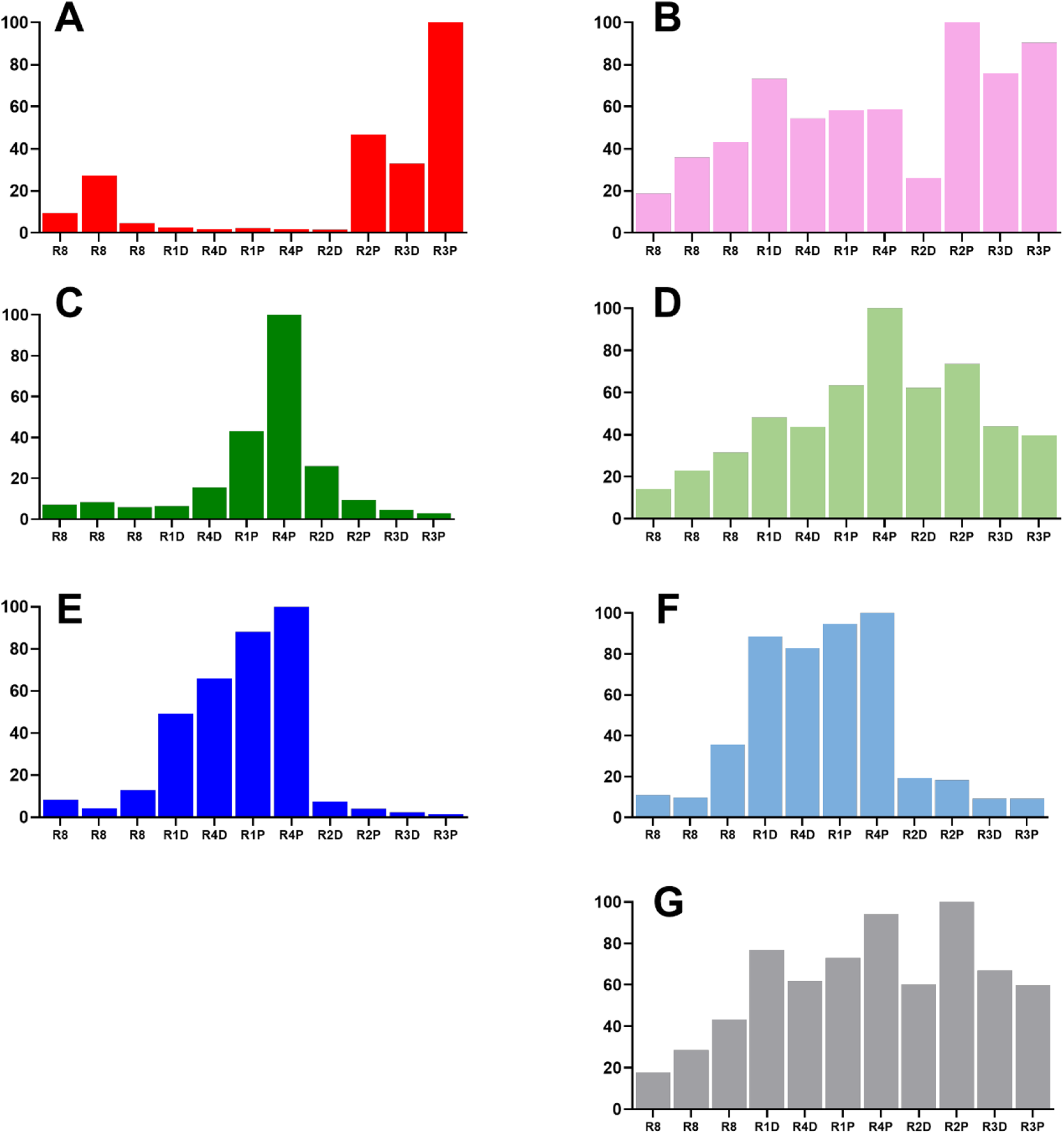
Photoreceptor responses to each filter type. Calculated relative quantum catch of each photoreceptor type to each coloured filter. **A)** highly saturated red **B)** less saturated red **C)** highly saturated green **D)** less saturated green **E)** highly saturated blue **F)** less saturated blue (the same used in Marshall et al., 1996) **G)** neutral density 0.15. Horizontal axis gives photoreceptor types (see **Fig. 1** for types).

### Experimental Procedure

Feeding choice tests were used to compare the ability of *H. trispinosa* to discern either high or low saturation colours from neutral greys. Each stomatopod was assigned a single colour high or low saturation red, orange, green, or blue. Animals readily emerge from their burrow to pick up the cable tie and retreat back into their home to consume the food (**Fig. 4**). A successful choice was recorded if they took the correct cable tie first and pulled it toward their burrow.

**Figure 4:**
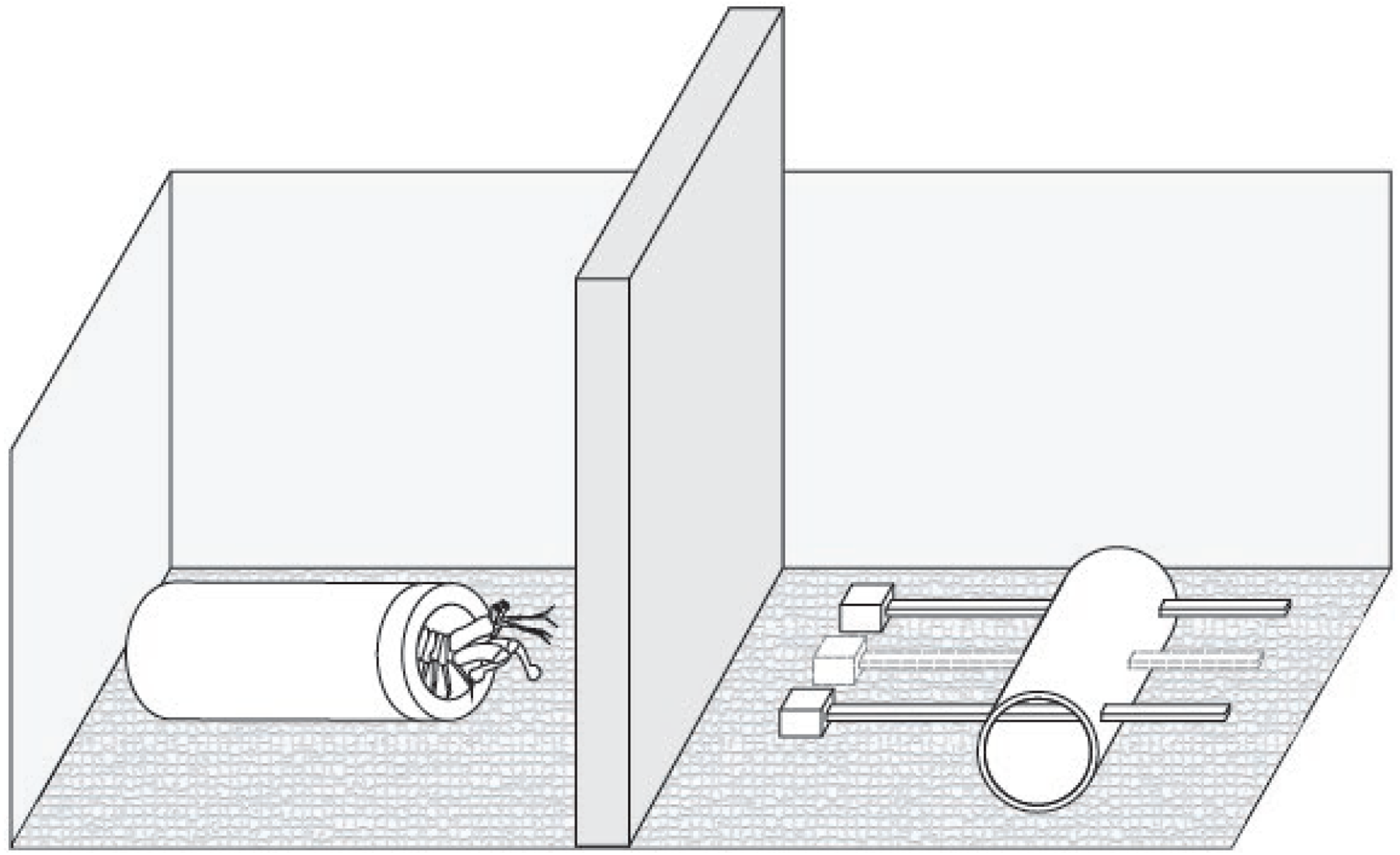
Behaviour setup. Setup just before an experiment begins. Stomatopod is in its burrow when a barrier is inserted and the cable ties placed (either two or three, see methods). The removal of the barrier signals the start of a trial, and the stomatopod will pull one cable tie into its burrow.

Animals were introduced to their stimulus during a “priming” week, where they were given a single cable tie, coloured according to their assignment, with food in the cavity created by the front of the cable tie. During the second week, they were “trained” with a priming cable tie (containing food), plus the distractor(s) (ND 0, 0.15, 0.6, 0.9) presented in a pseudo-random order. After the second week, the animals which reached approximately a 70% rate of cable-tie feeding were used for testing. Animals that did not participate were considered untrainable and were removed. This method is further described in Templin, 2017.

**Fig. 4** shows the experimental setup. During a trial, each stomatopod was presented with their assigned colour and one (Experiment 2) or two (Experiment 1) neutral density distractors. Cable ties were held loosely in a holder so they could be easily removed. The cable ties were presented approximately 3 cm from the entrance to their burrow as a set. In training trials, a small piece of food was placed inside the cavity of the target cable tie. During tests, the target cable tie and all neutral density distractors were newly fabricated and had never come into contact with food in order to eliminate any residual olfactory cues.

Tests in which cable ties were not in contact with food were conducted once a day, five times a week. Additionally, mantis shrimp were trained once a week with food present to reinforce the behaviour. The stomatopods were given approximately three minutes to make a decision or were considered to not have participated. A trial began when the barrier inserted between the burrow and the stimulus (**Fig. 4**) was lifted. If the mantis shrimp chose correctly, it was rewarded with a small piece of food. If it chose the unrewarded colour, the cable ties were removed, and no food was awarded.

The combination of coloured cable ties and neutral density filters was made using a random number generator. Enough experiments were conducted so that each combination occurred at least once during all testing trials but not more than three times during both training and testing. The trials were ordered such that the target cable tie was not in the same location for two (for 3-choice, Experiment 1) or three (for 2-choice, Experiment 2) consecutive trials. If a mantis shrimp participated 2 times or less per week, for more than one week, it was considered untrainable, and therefore replaced.

### Individual Experiments

#### Experiment 1: High/Low Saturation

In Experiment 1, we investigated the original “barcode” hypothesis, the possible expectation being that they would fail or find more difficult all low-saturation colours. Animals were tested across 8 colour types (four hues: red, orange, green, and blue, each at two saturation levels: high and low). Stomatopods were given a three-choice test, with two neutral density distractors. However, after a while it was noted that animals significantly preferred the middle position of the target cable tie (p < 0.001). Only one distractor was used in later experiments to prevent this bias, and in three-way choices, results were adjusted to account for this bias. These tests were exclusively conducted in captivity at UQ, beginning at least 3 months post-capture.

#### Experiment 2: High saturation discrimination over time

In the second experiment, we tested whether the length of time in captivity affected the ability of the mantis shrimp to discriminate colour. Tests consisted only of highly saturated red, green, and blue versus one neutral density distractor. Tests were divided into three time points to judge abilities at different lengths of time spent in captivity: (1) One week at LIRS; (2) weeks 1-10, at UQ under artificial lighting; and (3) at the final weeks of testing (weeks 10-20) at UQ. Animals at LIRS were trained and tested for one week, three times a day.

#### Experiment 3: Naïve-Choice Tests

All naïve-choice tests were conducted in the LIRS aquarium system under natural lighting in which freshly caught animals were given 24 hours to acclimate. Two-choice tests were conducted across all colours (red, orange, green, blue): (A) high saturation preference, (B) low saturation preference, (C) preference of colour versus neutral density grey, and (D) preference of high or low saturation. In high and low saturation tests (A and B), individuals were given a random pair of red, orange, green, or blue stimuli in the same saturation type. In the saturation preference test (C), individuals were given a high and low saturation stimulus of a single colour.

Finally, for colour versus grey preference (D), animals were given a randomised colour (red, orange, green, or blue) and neutral grey (ND 0, 0.15, 0.6, and 0.9) pair. Each individual was only used once in each experiment type; some were used in two different experiments due to collection restrictions. Individuals were given 30 minutes to make a choice. If an individual did not participate, it was tested again with the same choice for up to two additional sessions, and then replaced.

### Data Analysis

Ability to learn the task was evaluated with a simple binomial test (BINOM.DIST, Microsoft Excel). This approach compares the number of times they chose a particular stimulus with the number of times they would be expected to choose it by chance. Individuals were included in analysis only if they were successful more often than expected by chance during training trials, as well as those which participated in at least 10 trials in Experiment 1, at least 3 trials at LIRS, or 5 trials at UQ in Experiment 2. Animals were assumed to have learned the task if they selected the target stimulus more often than chance. Experiment 3 analysis was also performed with a binomial test to determine significance between the two choices.

Data from Experiments 1 and 2 were analysed using the general linear mixed effects model package in R (lme4 package, R version 3.5.3; Bates, Martin and Bolker, 2011). Success for each experiment was used for the response variable in all analysis. Depending on the experiment, the colour (red, orange, green, blue), colour type (high or low saturation), time period, and neutral density distractors were treated as fixed factors. The individual ID of each animal was treated as a random factor in all analyses.

To understand interacting effects, including the effect of the neutral density filter types and cable tie position, a post-hoc test was performed for individual colours and colour types using Tukey’s multiple comparisons test (Hothorn et al., 2015). Only data from “test” trials were used, and only when the stomatopod performed the task.

## Results

### Experiment 1: High/Low Saturation

*H. trispinosa* were able to learn to distinguish all colours from grey (binomial test: highly saturated orange p < 0.05 and highly saturated blue p < 0.01, p < 0.0001 for all others). There was an overall trend towards better performance in less saturated colours (**Fig. 6A**). This was significant for red (GLMM, p < 0.05) and blue (p < 0.0001), but not orange (p > 0.05) or green (p > 0.1) (**Table 1**).

**Table 1:**
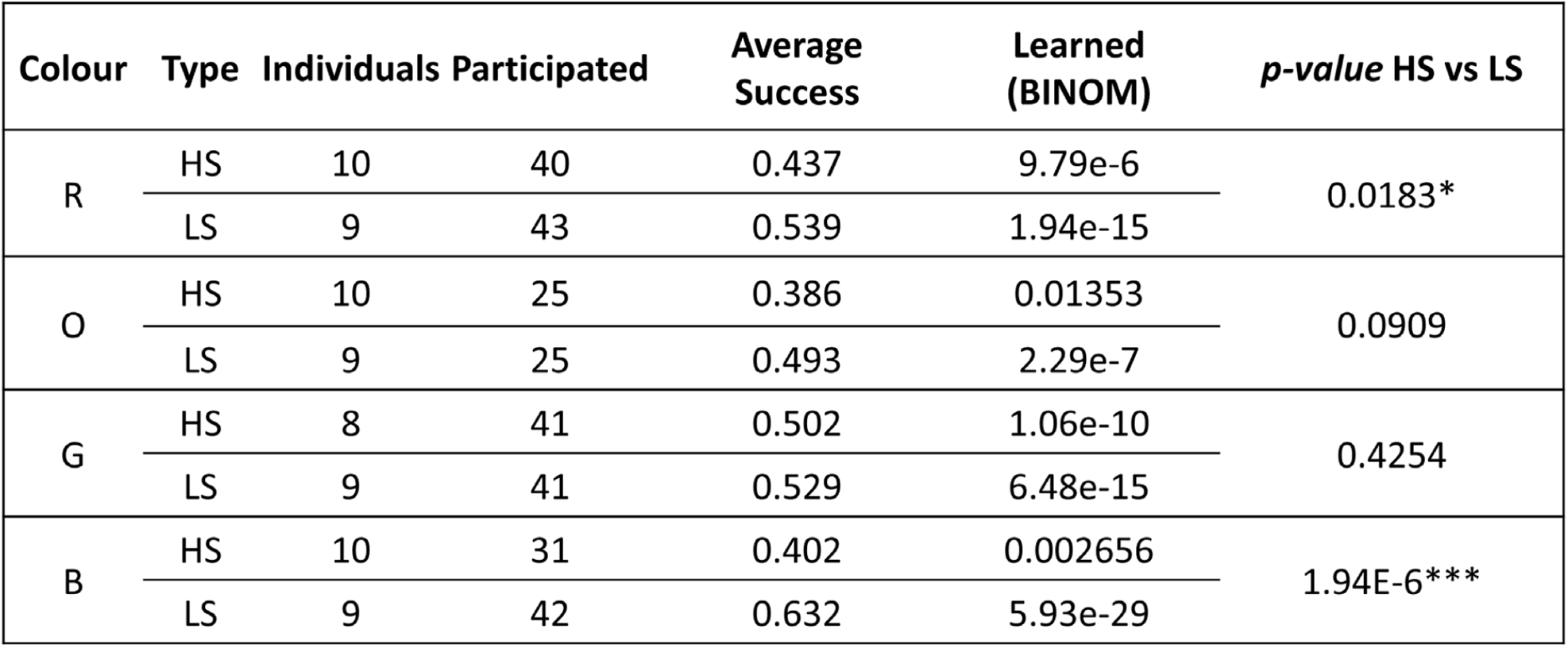
High and low saturation learning. Number of individuals, average number of trials, average success, learning ability, and difference between high (HS) and low (LS) saturation for each colour and colour type. The trials included 3 choices, suggesting that an average success rate over 0.333 corresponds to learning the task.

**Figure 5:**
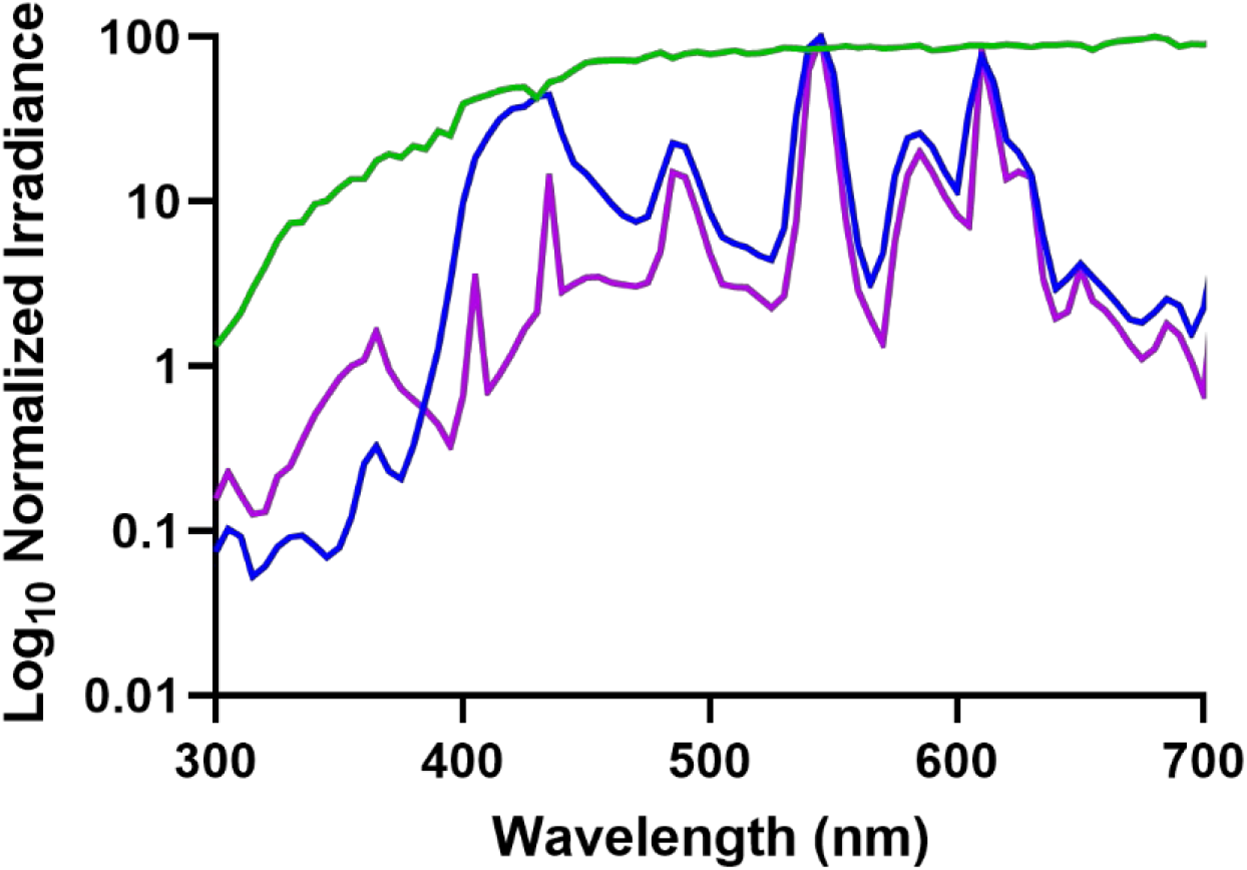
Available light spectra. Available light in natural environment at LIRS (green) and overhead lights in UQ Lab: Experiment 1 (blue) and Experiment 2 (purple). Irradiance is normalized for each measurement to show relative spectral availability.

**Figure 6:**
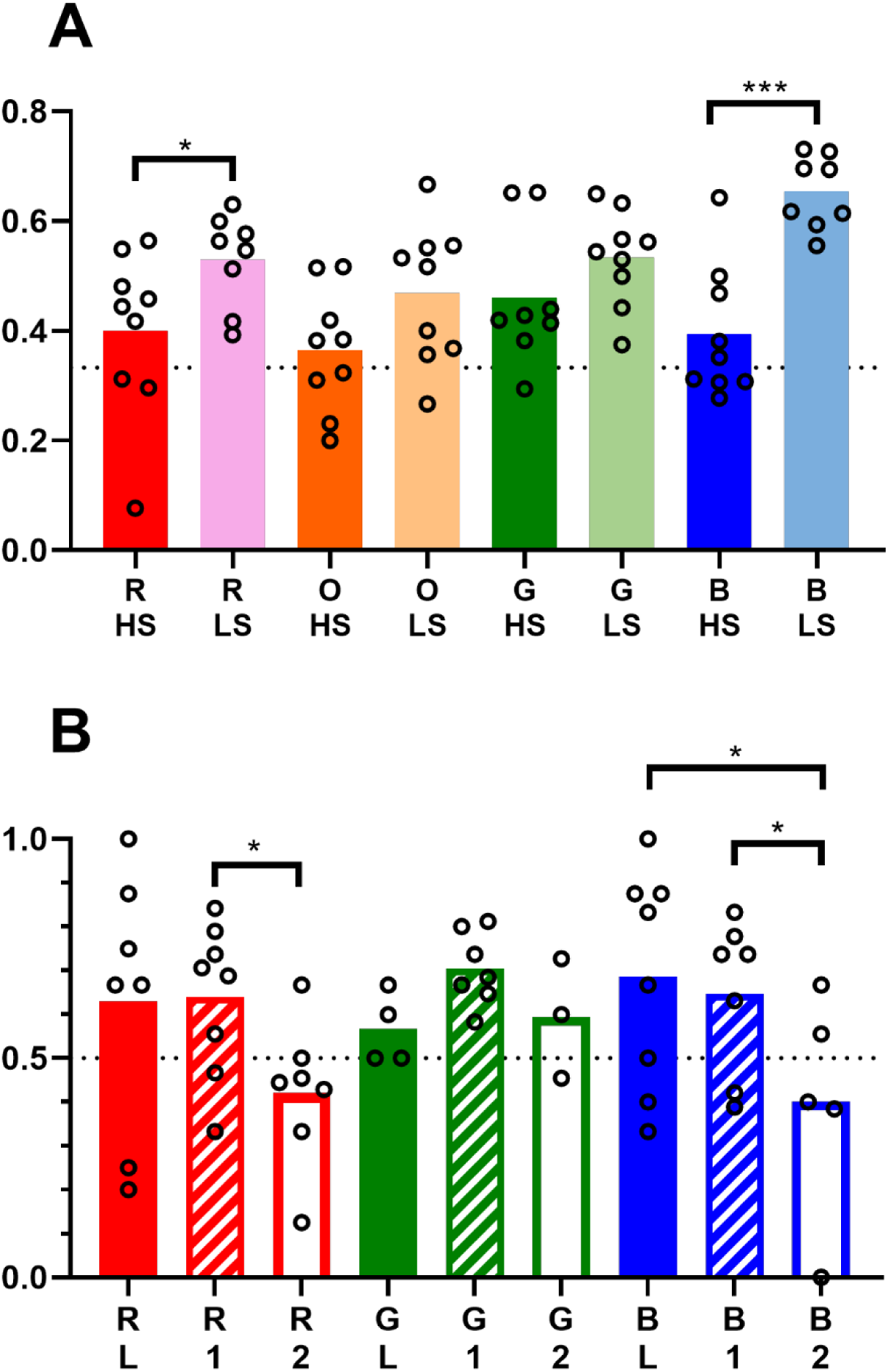
*H. trispinosa* performance in captivity. Average performance of individuals in each category. **A)** Experiment 1 results, comparison of high saturation (HS) and low saturation (LS) stimuli. **B)** Experiment 2 results, performance over time in captivity, at LIRS (“L”), first 10 weeks (“1”), and second 10 weeks (“2”). Dotted black line indicates the level of chance. Circles indicate each individual’s average.

An average of 9±1 individuals per colour type participated in an average of 36 trials (range 10-71, depending on trainability and mortality). There was no significant variation between individuals (model variance < 0.1). Overall, many animals were less successful on trials when at least one of the distractors was ND 0.15 and/or 0.6, especially in less saturated red and highly saturated green (**Fig. S1.1, Table S1.1**). These neutral density distractors were similar in brightness to the target stimuli (**Fig. 2**). There was no significant effect of sex or size on the overall success, therefore these factors were not included in further analysis (p > 0.5).

### Experiment 2: Discrimination over time

*H. trispinosa* were trained to discern red, green, and blue high saturation stimuli from a neutral grey distractor, at LIRS and UQ. Over the first set of ten weeks in captivity, all animals were able to learn the task successfully (p < 0.001). In the second set of ten weeks, accuracy was severely diminished. During this time, animals were not able to choose the coloured stimulus significantly more than chance for all colours (p > 0.05) (**Table 2**).

**Table 2:** Discrimination over time. **A, B)** show *p-values* of mixed models using GLMM (success ∼ set + colour + ND + [1|individual]) and further Tukey tests on the factors. **A)** Effect of set, by colour. **B)** Effect of colour, by set. **C)** Number of individuals and average participation per colour per set, along with learning performance (see methods). Average success is in a 2-choice trial, and thus a success rate of 0.5 would correspond to chance.

| **A** | 1 vs 2 | 1 vs 3 | 2 vs 3 |
| --- | --- | --- | --- |
| All | 0.9482 | 0.0288* | <0.001*** |
| R | 0.9858 | 0.2449 | 0.0368* |
| G | 0.524 | 0.999 | 0.374 |
| B | 0.9240 | 0.0431* | 0.0207* |

| **B** | R v G | R v B | G v B |
| --- | --- | --- | --- |
| All | 0.461 | 0.899 | 0.722 |
| Set 1 | 0.966 | 0.748 | 0.640 |
| Set 2 | 0.736 | 0.994 | 0.685 |
| Set 3 | 0.273 | 1.000 | 0.314 |

**Table 2: Discrimination over time.**
| **C** | Colour | Individuals | Participated | Average Success | Learning (BINOM) |
| --- | --- | --- | --- | --- | --- |
| Set 1 | R | 7 | 4.3 | 0.633 | 0.0509 |
|  | G | 4 | 4.75 | 0.579 | 0.1442 |
|  | B | 8 | 5.6 | 0.717 | 0.00145** |
| Set 2 | R | 8 | 17.25 | 0.652 | 1.08e-4*** |
|  | G | 7 | 16.6 | 0.704 | 2.91e-6*** |
|  | B | 7 | 18.6 | 0.646 | 2.61e-4*** |
| Set 3 | R | 7 | 8.7 | 0.443 | 0.0685 |
|  | G | 3 | 12.3 | 0.595 | 0.0681 |
|  | B | 5 | 9 | 0.444 | 0.0901 |

For all colours, *H. trispinosa* became significantly less successful at selecting the correct stimuli after spending time in artificial light (GLMM p < 0.001). After the first 10 weeks in captivity, animals became less effective at selecting red and blue high saturation stimuli (p < 0.05). In addition, animals learning highly saturated blue were more successful at LIRS in natural light than in the artificial lab light (p < 0.05) (**Fig. 6B, Table S1.2**).

Individual variance in the linear mixed model was less than 0.2, and lower for most colours and sets, except for highly saturated blue and at LIRS (approximately 0.5). On average, 6 individuals were used per colour with an average of 5 trials each at LIRS; 7 individuals per colour with 17 trials each during the first ten weeks at UQ; and 5 individuals with 10 trials each during the second ten weeks at UQ (**Table 2**). Animals did significantly better in trials with neutral density filter ND 0 in all trials (GLMM p < 0.0001), when trained to green and blue (p < 0.01 and p < 0.05, respectively) and on LIRS and the first 10 weeks at UQ (p < 0.05 and p < 0.0001, respectively) (**Table S1.2**).

### Experiment 3: Naïve-Choice Tests

*H. trispinosa* were given a two-choice test to determine if there was an innate preference for any of the colours or colour types. There was a significant preference for red (binomial test, p < 0.01) and aversion to green (p < 0.05) among highly saturated colours (**Fig. 7A**). No preference was found for any low saturation colours (**Fig. 7B, Table S2.1**).

**Figure 7:**
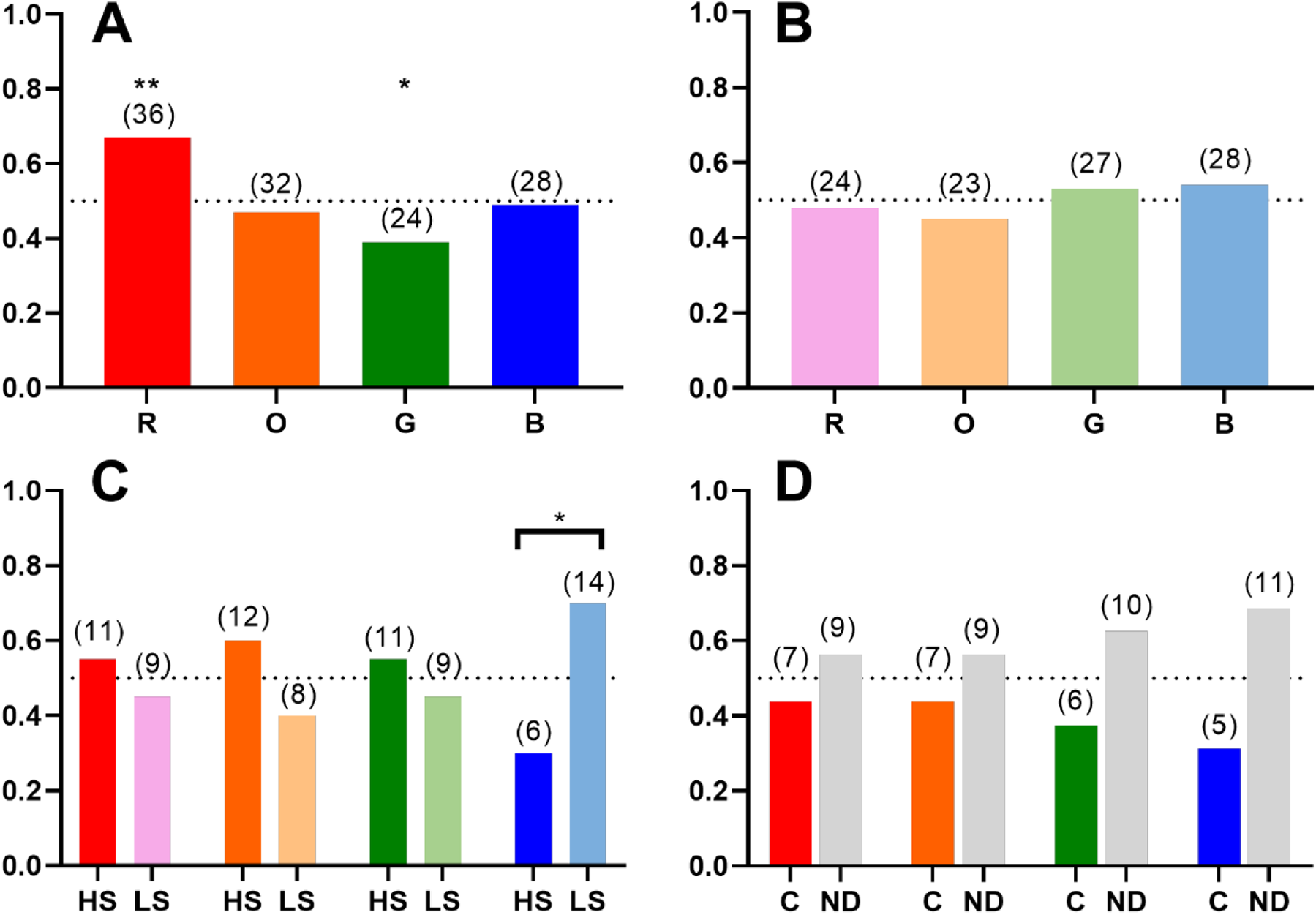
H. trispinosa naïve-choice tests. **A, B**) Results from a two-choice test between each colour in **A**) high saturation and **B**) low saturation colours (i.e. R/O, R/G, R/B, O/G, O/B, G/B). **C**) Choice between high saturation (HS) and low saturation (LS) stimuli of the same colour. **D**) Choice between a high saturation colour (R, O, G, B) and neutral-density grey (ND 0, 0.15, 0.6, 0.9). Black line refers to the level of chance. Numbers refer to number of individuals with the respective preference.

Other individuals were given a choice between the high and low saturation stimuli of each colour. Animals preferred low saturation to high saturation blue (p < 0.05) but had no preference between saturation types for the other colours (**Fig. 7B, Table S2.1**). When animals were given a choice between a colour filter and neutral density filter, stomatopods preferred the neutral density to all colours (p < 0.05), but this was not significant when colours were analysed separately (**Fig. 7C**). However, they significantly avoided the brightest neutral density filter (ND 0, p < 0.05) (**Fig. S1.2B**).

## Discussion

The aim of this study was to examine colour processing in *H. trispinosa*, by exploring the two current hypotheses, opponent processing and pattern activation (also called barcode or cochlea-vision). Our initial hypotheses were that stomatopods may: a) fail at all low-saturations colours (mirroring and extending the results of Marshall et al., 1996 and Thoen et al., 2014) or b) that they may just fail with low-saturation blue, suggesting some behavioural relevance or significance for this colour. This would have provided evidence, albeit circumstantial, that their colour processing was somehow different and potentially bar-code-like. It should be noted that Marshall et al. (1996) did not exclude opponent processing for colour vision, pointing out that the chromatic signal differential was lowest for the low-saturation blue which that species (*O. scyllarus*) failed on.

Results of our study did not clarify this debate in stomatopod vision (Marshall and Arikawa, 2014; Marshall et al., 1996, 1997; Thoen et al., 2014). Interestingly, some of our findings raise questions regarding the methodologies in previous and current experiments regarding animal colour vision capabilities. Hence the reason for Experiments 2 and 3 conducted here as discussed below.

We found that *H. trispinosa* could learn to discriminate both high and low saturation colours from grey. This was an unexpected result as we originally proposed all low saturation colours would be harder to discriminate based on the results of Marshall et al., 1996 and Thoen et al., 2014 (**Fig. 6A**). H. *trispinosa* demonstrated relatively poor discrimination of high saturation orange and blue colours, compared to other colours, however we suggest this was due to length of time spent in captivity, rather than an accurate reflection of their visual capabilities. Extended time in captivity may reduce visual capacity due to the light spectrum being more limited than natural daylight, or possibly other influences such as restricted diet (**Fig. 6B**). Most notably, stomatopod require carotenoids for the formation of red filters in the retina, and these must be obtained through their diet (Cronin and Caldwell, 2002; Marshall et al, 1991b). In light of this, the previous finding that *O. scyllarus* could not discriminate low saturation blue may be a product of spending time in captivity. In particular, due to the assumption that if humans can see the colour, another animal with apparently superior colour vision must be able to, little effort was made to imitate natural daylight. Animals were kept in an inside room with no window and whatever fluorescent strip-light present in the room at the time. Furthermore, as animals were sourced from a tropical marine supplier, the amount of time in captivity prior to experiments starting was unknown. An alternative explanation for the apparently contradictory results we obtained is one of species specificity. Blue and indeed other low-saturation colours may be more salient and important for *H. trispinosa* than *O. scyllarus*.

Since 1996, it has been demonstrated that the spectral sensitivities of stomatopods, and indeed other animals, are remarkably plastic, being influenced by both depth and light environment, as well as changing seasonally or on even shorter timescales, such as days or within a single day (crustaceans: Cronin et al., 2002; Jessop et al., 2020; fishes review: Carleton et al., 2020; Musilova et al., 2021). Our results with *H. trispinosa* further support this visual plasticity, and suggest it is possible, that the animals trained in 1996 had a short-term-modified colour sense significantly different to the one ordinarily present under natural lighting conditions (**Fig. 6**). Even in the wild, stomatopods are known to modify and tune spectral sensitivities depending on the spectral envelope of ambient light they live in, according to the depth of habitat they settle into as post-larvae (Cronin and Caldwell, 2002; Cronin et al., 2000, 2001; Cheroske et al., 2006; Osorio et al., 1997).

There are other historical problems with the study of Marshall et al. (1996). For example, at that stage the relative sensitivity of each row based on optics were unknown, thus non-normalised spectral sensitivities were used to calculate and compare quantum catch between photoreceptors (Cronin et al., 2000). This resulted in an apparent closer match between quantum catch pattern looking at ND filters and the unsaturated blue (Marshall et al., 1996 Fig. 4). Quantum catch calculations performed here (**Figs 2F, G**) where the same blue filter was used but with more uniform absolute sensitivities in each row, may be more accurate.

The results of Experiment 1, led to the conception of Experiment 2, to determine if the previous findings of Marshall et al (1996) could be explained by time in captivity and/or lighting conditions. Interestingly, stomatopods had trouble distinguishing highly saturated red and blue after three months in captivity (**Fig. 6B**), supporting the idea that the negative result of Marshall et al. (1996) needs to be viewed with caution or may indeed be wrong. More recently we have discovered that mantis shrimps are able to shift their spectral sensitivity under different light environments, both natural and unnatural, and do so on the same timescales seen here (Cheroske et al., 2003, 2006; Cronin et al., 2000). This shift is usually towards shorter wavelengths in the more red-sensitive row 3, an adaptative response to the reduced long wavelengths in deeper or bluer habitats (Cheroske et al., 2006; Cronin et al., 2002; Cronin and Caldwell, 2002). The result here suggests that the stomatopods used in Marshall et al. (1996) had a change in colour discrimination ability and that the failure to discriminate unsaturated blue from grey may have been a result of this short-term adaptation process.

In response to the results from Experiment 1 - where *H. trispinosa* displayed a reduced ability to learn saturated orange and blue - we also conducted Experiment 3 to test if *H. trispinosa* has an innate avoidance or preference for colours used in this experiment. *H. trispinosa* display bright blue markings on the carapace and maxillipeds in aggressive and mating contexts (Chiou et al., 2005) making it possible that blue may have attached to it a specific significance. These and other species have been found to spontaneously avoid UV markings, also a feature of the frontal displays of stomatopods when they meet or contest ownership of cavities within which to live on the reef (Bok et al., 2018). Our results indicate that *H. trispinosa* did not show any specific avoidance of highly saturated blue but did display a preference for red and an avoidance of green (**Figs 7A, S2.1A**). The preference for longer wavelengths is similar to the results from Daly et al. (2017) where a colour preference for yellow and an avoidance of red was found in naïve choice tests in the peacock mantis shrimp, *O. scyllarus*. The nature of the tests conducted was different, but this comparison highlights a potential difference in colour behaviour between species. In addition, we found that *H. trispinosa* had no preference among low saturation colours (**Fig. 7B**), but did prefer low over high saturation blue (**Fig. 7C**). This may account for the difference in success rate between high and low saturation blue in the first experiment and may indicate an underlying avoidance of blue in this species (**Figure 6A**)

Different specific colour preference in naïve choice tests have been found in other crustaceans. For example, male blue crabs, *Callinectes sapidus*, show a preference for red over orange claws in females, as mature females have red claws while those of prepubertal females are orange (Baldwin and Johnsen, 2012). On the other end of the spectrum, fiddler crab (*Uca mjoebergi*) females prefer males whose yellow claws are also UV reflective (Detto and Backwell, 2009). Among the insects, innate long-wavelength preference also occurs in some species of butterflies (Kinoshita and Arikawa, 2014; Swihart and Swihart, 1970; Weiss, 1997) and flower-pollinating flies (An et al., 2018; Lunau et al., 2018).

Interestingly, naïve-choice tests showed that *H. trispinosa* had a preference for medium-brightness grey over high saturation colours (**Figure 7D**). Conversely, innate preference tests in crabs show that they prefer colours over grey, which is suggested due to use of colour in sexual selection. Female fiddler crabs prefer yellow over grey, similar to male claw colouration (Detto, 2007), while male blue crabs prefer the red claws, which mature females exhibit, over white and black claws (Baldwin and Johnsen, 2009).

To find if the ability to distinguish both low and high saturated colours is more widespread among stomatopods, we repeated experiments 1 and 2 with *G. smithii*. In common with *H. trispinosa*, there was a trend towards learning low saturation colours, and degradation of results over time kept in captivity (**Fig. S3.1, Table S3.1**).

Although the set of experiments described here have further explored the colour discrimination of stomatopods, even the basics of the colour processing mechanism remain unclear. It is possible that stomatopods use a combination of multiple mechanisms to process colour information in different behavioural contexts, including opponency and photoreceptor activation comparisons, or bar-code analysis. In addition, although the retinal design and underlying structures are largely similar in all mantis shrimp species with six-row midbands, different species may process colour differently (Marshall et al., 2007; Thoen et al., 2017).

Given that mantis shrimps display species-specific colour markings during encounters between and within species on the reef (Caldwell and Dingle, 1975) it is worth considering if this aspect of behaviour plays a stronger part in stomatopod colour vision and that the colour of food, or food containers, is irrelevant in normal life. During aggression sequences, often both mantis shrimp spread their front raptors to show the species-specific colour of their meral spot to evaluate their opponent; an act which may either lead to fighting or submission (Caldwell and Dingle, 1975; Dingle and Caldwell, 1969; Green and Patek, 2015, 2018). Both intensity and chromaticity of the meral spot is a signal of aggression in some species: a darker meral spot indicates a stronger strike force, and a lighter meral spot often leads to the receiver increasing antagonism (Caldwell and Dingle, 1975; Franklin et al., 2017, 2019; Hazlett, 1979).

While the results of experiments here have not led to firm conclusions, what is clear is that a number of previous experimental protocols in colour vision experimentation may need adjusting. The fact that vision changes over evolutionarily short time spans (reviewed in: Land and Nilsson, 2012; Nilsson, 2013; Kelber and Osorio, 2010; Marshall et al., 2015) and apparently colour vision is remarkably plastic in both vertebrates (Carleton et al., 2020; Kelber et al., 2003; Musilova et al., 2021) and invertebrates (Cronin et al., 2002; Jessop et al., 2020; Strausfeld and Andrew, 2011; van der Kooi et al., 2020) on very short time scales presents a fascinating area for further study. Does a change in visual pigment expression level or photoreceptor complement lead to a change in colour detection or discrimination? What degree of colour constancy underlies these changes? Is a food reward-based behavioural experiment sufficiently basal that other colour-based behaviours, such as mate choice or aggressive interaction, simply follow suit; or are there different levels of discrimination for different behavioural tasks? As usual with the stomatopods, we have found more questions than answers.

## Acknowledgements

Thank you to Dr. Karen Cheney for assistance in interpreting mixed model results. Thank you also to Connor Brainard, Justine Ohlrich, and Lena van Swinderen for help in performing experiment 1 trials.

## Author Contributions

AS and JM conceived and designed the experiment; AS and HE collected data; AS analysed data and AS and JM wrote the manuscript; all authors edited the manuscript.

